# Effects of water content and mesh size on tea bag decomposition

**DOI:** 10.1101/2020.11.15.384016

**Authors:** Taiki Mori, Ryota Aoyagi, Hiroki Taga, Yoshimi Sakai

## Abstract

The tea bag method was developed to provide uniform litter bags that enable comparison of organic matter decomposition rates on a large scale. However, it remains uncertain whether tea bag decomposition in response to wetness is representative of that of natural litters. We performed incubation experiments to examine whether the effect of soil water on tea bag decomposition becomes inhibitory at higher water contents, as was demonstrated in natural leaf litters. In addition, we performed field studies in a mixed forest and cedar plantation in Japan to compare two litter bag mesh sizes: 0.25-mm mesh, the size previously used by a major manufacturer of tea bags (Lipton), and nonwoven bags with mesh sizes finer than 0.25 mm, which are currently produced by Lipton. Both green tea and rooibos tea exhibited higher decomposition rates at higher water contents, but decomposition was inhibited at the highest water content, consistent with conceptual models of natural litters. The nonwoven tea bags did not show lower decomposition rates, despite the finer mesh size. Rather, the nonwoven rooibos tea bags exhibited slightly higher decomposition rates than the 0.25-mm mesh bags in the cedar plantation, possibly due to a greater abundance of microorganisms that decompose litters in the nonwoven bags, due to the decrease in predation by mesofauna. Our findings provide essential information for future studies of tea bag decomposition.

## Introduction

Recent anthropogenic activities have caused global issues such as climate change and nitrogen loading, which may have a considerable negative impact on biodiversity and ecosystem services. To understand the long-term impact of these global issues on ecosystems, it is essential to examine how environmental perturbation at a large scale affects litter decomposition rates, because decomposition is a basic ecosystem process required to sustain nutrient cycling. However, the commonly used litter bag method has difficulty in detecting the effects of environmental factors on decomposition at a large geographical scale because it is difficult to standardize litter quality, which is a primary factor controlling litter decomposition rates (Cornwell et al. 2008, Zhang et al. 2008).

The tea bag method, introduced by Keuskamp et al. (2013), provides standardized litter decomposition data due to uniformity of the litters and bags, which is necessary for large-scale analysis of the effects of environmental factors on decomposition rates. The tea bag method uses commercially available tetra-shaped tea bags of green tea (*Camellia sinensis*) and rooibos tea (*Aspalathus linearis*), produced by Lipton, as a substitute for litters and soil organic matter. The amount of tea leaves decomposed during a 90-day incubation period is used to calculate the decomposition constant *k* and stabilization factor *S* (Tea Bag Index) (Keuskamp et al. 2013). Temporal variations in chemical composition during the decomposition process exhibited were typical of natural leaf litters (Duddigan et al. 2020), indicating that tea bags can be representative of natural litters. A large number of studies have used the tea bag method to assess decomposition potential (Fujii et al. 2017, Djukic et al. 2018, Mueller et al. 2018, Petraglia et al. 2019, Suzuki et al. 2019).

However, it remains uncertain whether tea bag decomposition in response to wetness is representative of that of natural litters. Generally, higher decomposition rates are observed in wetter conditions, but excess water inhibits decomposition: according to Prescott (2010), water contents higher than 80% (wet weight basis) suppresses litter decomposition rates. By contrast, a field experiment reported that tea bag decomposition was not suppressed, even at high water contents, based on comparison of tea bag decomposition data obtained from various study sites (Petraglia et al. 2019). Tea leaves may have large water-leachable fractions, where the inhibitory impact of very high water content on litter decomposition may be negated by large leaching fractions under wetter conditions. If this is the case, tea bags should not be used to determine litter decomposition rates under high soil water contents. In field studies, however, other environmental factors covary with soil water content, so it remains unclear whether the effect of soil water on tea bag decomposition becomes inhibitory at higher water contents. In the present study, we performed laboratory incubation and leaching experiments to clarify the effect of water content on tea bag decomposition, and the potential contribution of leaching loss.

To utilize the tea bag method to test the impact of environmental changes on litter decomposition at a large scale, a practical issue must be resolved first. Lipton changed the mesh materials for its tea bags from woven nylon mesh (0.25 mm) to polypropylene nonwoven mesh (non-uniform size, but finer than 0.25 mm; see Fig. S1) in 2017. Since the previously used woven bags are unavailable, future works should use the new nonwoven bags. However, we are uncertain whether the newly obtained data can be combined with previous data obtained using the 0.25-mm mesh bags. Because mesh size generally affects the accessibility of decomposers and microclimate in litter bags (Bradford et al. 2002, Powers et al. 2009), any change in mesh size of tea bags is likely to affect the decomposition rate. To date, no data have shown the impact of mesh size (0.25-mm mesh vs. nonwoven mesh) on tea bag decomposition. In the present study, we compared two types of tea bags with different mesh sizes, namely, 0.25-mm mesh bags and nonwoven bags with mesh sizes finer than 0.25 mm. We predicted that the rooibos tea decomposition rate data obtained from the two types of bags would not be amenable to being combined, because the finer mesh size would lead to a lower decomposition rate of rooibos tea, as the finer mesh causes slower decomposition (Bradford et al. 2002, Powers et al. 2009). On the other hand, the decomposition rate of green tea may not be affected by the difference in mesh size, given that (i) easily decomposable fractions of green tea are typically completely decomposed before the end of the 90-day incubation period (Keuskamp et al. 2013) and (ii) recalcitrant fractions are not decomposed by microbes or mesofauna during the 90-day incubation (Keuskamp et al. 2013).

## Materials and methods

### Study sites

The experiments were performed in a mixed forest dominated by *Chamaecyparis obtusa* and *Clethra barbinervis*, and in a Japanese cedar (*Cryptomeria japonica*) plantation. The mixed forest (35.07 N, 135.76 E) is located at the Kamigamo experimental station in Kyoto, Japan. The cedar plantation (32.82 N, 130.73 E) is located at the Tatsudayama research site in Kumamoto, Japan. The mean annual temperature and precipitation amounts at the research sites were 15.6 and 1,932 mm, respectively, in the mixed forest, and 17.1°C and 1,951 mm, respectively, in the cedar plantation. The climate data were obtained from The Agro-Meteorological Grid Square Data of the National Agriculture and Food Research Organization (NARO).

### Tea bags

Following the developers of the tea bag method (Keuskamp et al. 2013) and their webpage (Teatime 4 Science; http://www.teatime4science.org/), green tea bags (EAN: 87 10908 90359 5; Lipton) and rooibos tea bags (EAN: 87 22700 18843 8; Lipton) were used. The current tea bags (polypropylene nonwoven tea bags) were used for the laboratory studies testing the impact of water content on tea bag decomposition. To examine the effect of mesh size on tea bag decomposition, we produced 0.25-mm mesh woven tea bags from the tea contained within the nonwoven tea bags and a 0.25-mm mesh (Fig. S2). Several chemical analyses were done in previous studies. According to Keuskamp et al. (2013), the fractions of nonpolar extractives, water solubles, acid solubles, and acid insolubles were 0.066 ± 0.003, 0.493 ± 0.021, 0.283 ± 0.017, and 0.156 ± 0.009, respectively, in green tea, and 0.049 ± 0.013, 0.215 ± 0.009, 0.289 ± 0.040, and 0.444 ± 0.040, respectively, in rooibos tea. Total C and N contents were 49.055 ± 0.109% and 4.019 ± 0.049%, respectively, in green tea, and 50.511 ± 0.286% and 1.185 ± 0.048%, respectively, in rooibos tea (Keuskamp et al. 2013). Duddigan et al. (2020) provided chemical composition data, obtained by nuclear magnetic resonance. Fractions of alkyl C, O-alkyl C, aromatic C, and carbonyl C were 0.230 ± 0.032, 0.570 ± 0.003, 0.146 ± 0.020, and 0.054 ± 0.009, respectively, in green tea, and 0.152 ± 0.044, 0.714 ± 0.018, 0.102 ± 0.017, and 0.032 ± 0.009, respectively, in rooibos tea.

### Laboratory studies

At the Tatsudayama research site, two incubation experiments and two leaching experiments were conducted to clarify (i) the effects of water content on the tea bag decomposition rate and (ii) the potential contribution of leaching loss to the rate. First, a laboratory incubation experiment was done to test the effects of soil water content on tea bag decomposition (Water Content Experiment). Second, the potential contribution of leaching loss to the tea bag decomposition rate was estimated by calculating the ratio of the leached amount to the mass loss. The maximum and minimum leaching losses were estimated by two leaching experiments (Maximum Leaching Experiment and Minimum Leaching Experiment). The mass loss was estimated by another incubation experiment, which involved treatment with a sufficient amount of water (Total Mass Loss Experiment). In the present study, we used the relative mass loss amount of teas (i.e., loss weight / initial weight) during a certain interval as an indicator of the leaching or decomposition rate, rather than the decomposition constant *k*.

### Water Content Experiment

In October 2019, soil samples (0–10 cm depth) were taken from the Tatsudayama research site in Kumamoto, Japan. We sieved the soils through a 4-mm sieve after removing large pieces of organic matter. We placed 150 g of fresh soil and two tea bags (green and rooibos teas, nonwoven tea bags) in a polyethylene terephthalate (PET) bottle (without a drainage hole). Water contents were adjusted to 27%, 39%, 48%, and 56% by adding deionized water (wet weight basis). The bottles were incubated for 90 days at 25°C in the dark. Four replicates were prepared for each treatment. Dry weights of the tea bags were determined immediately after the end of the incubation.

### Total Mass Loss Experiment

*S*oil samples taken from the 0–5-cm depth in September 2019 were used for the Total Mass Loss Experiment. Soils were sieved through a 2-mm sieve after large pieces of organic matter had been removed. Fresh soil (70 g) and two nonwoven tea bags (one green tea and one rooibos tea) were placed into a PET bottle with a drainage hole at the bottom (see Fig. S3). The bottles were incubated for 90 days at 3°C (in a cold room) or 25°C (in an incubator). Four replicates were prepared for each treatment. To prevent water evaporation, all bottles were covered with a polyethylene sheet (Mori et al. 2013). At 10, 23, 57, and 90 days after the start of the incubation, 100, 100, 200, and 200 mL of deionized water was added to simulate precipitation, respectively. In several samples, the added deionized water (200 mL) on the last day (day 90) did not drain for several hours, so the incubation ended without complete drainage of the added water. Tea bags were oven-dried immediately after the end of the incubation period to prevent further decomposition. After removing the polypropylene fabric and soils on the surface of the fabric, teas were placed in an oven again (70°C for 72 h) and dry weights were determined.

### Minimum Leaching Experiment

A leaching experiment was conducted to determine the minimum leaching losses, using the same soils as in the Total Mass Loss Experiment described above. To exclude any impact of microorganisms on mass loss of the tea bags, soils were autoclaved at 120°C for 1 h. The autoclaved soils (70 g) and two nonwoven tea bags (green and rooibos teas) were placed into the PET bottle with a drainage hole (Fig. S3). Distilled water (600 mL) was added at 3°C (in a cold room) or 25°C (in an incubator). Three replicates were prepared for each treatment. After the water had been drained, tea bags were oven-dried immediately, and their dry weights were determined as described above.

### Maximum Leaching Experiment

The maximum leaching loss of the tea bags was measured based on a modified version of the method of Nykvist (1959). Tea bags (nonwoven tea bags) were submerged in 300 mL of deionized water in glass or plastic bottles, and leaching loss at different temperatures and submerging durations was determined. Leaching proceeded in a cold room (3°C) or incubators (15°C and 25°C) for 10 min, 140 min, or 24 h. Three replicates were prepared for each treatment. Dry weights were determined immediately after the end of the experiment.

### Field experiments

Tea bag decomposition rates were compared between 0.25-mm mesh tea bags and nonwoven tea bags in the mixed forest at Kamigamo research station and the Japanese cedar plantation at the Tatsudayama research site. At each study site, we used four sub plots. Pairs of 0.25-mm mesh tea bags and nonwoven tea bags (both green and rooibos tea bags) were buried with two replicates at 8-cm depth, following Keuskamp et al. (2013). The bags were re-collected after 90 days. Dry weights (oven-dried at 70°C for 72 h) were measured after removal of the mesh and soils on the surface thereof. The 90-day field incubation was initiated in October 2019 and September 2019 in the mixed forest and Japanese cedar plantation, respectively. The average temperature and cumulative precipitation during the 90-day field incubation experiments was 10.9°C and 116 mm, respectively, in the mixed forest, and 15.5°C and 359 mm, respectively, in the cedar plantation (Fig. S4, obtained from The Agro-Meteorological Grid Square Data, NARO).

### Calculation and statistics

The lower bounds of the potential contribution (minimum contribution) of leaching loss to the tea bag decomposition rate was estimated by calculating the ratio of the minimum leaching loss (determined in the Minimum Leaching Experiment) to the total mass loss (determined in the Total Mass Loss Experiment). The upper bound of the potential contribution (maximum contribution) of leaching loss to the tea bag decomposition rate was estimated by calculating the ratio of the leaching loss during 24-h submergence in water (determined in the Maximum Leaching Experiment) to the total mass loss (determined in the Total Mass Loss Experiment), assuming that the leachable fraction of the teas was mainly lost during the 24-h submergence period.

Statistical differences among treatments were tested using analysis of variance (ANOVA; one-, two-, or three-way) and the paired *t*-test, assuming a normal distribution of the data. Tukey’s HSD was used as a post hoc test as necessary. All statistical analyses were performed using R software (version 3.5.3 and 4.0.2; R Development Core Team 2019, 2020).

## Results

### Effects of water content on tea bag decomposition

The Water Content Experiment revealed the relationship between soil water content and tea bag decomposition rate. Two-way ANOVA showed that the interactive effect of moisture and tea type was significant (Fig. 1). According to the post hoc analysis, (i) up to a gravimetric soil water content of 48% (wet weight basis), the tea bag decomposition rate increased with increasing soil water content, but (ii) decomposition was suppressed at the highest water content in both green tea and rooibos tea (Fig. 1).

**Fig. 1.**
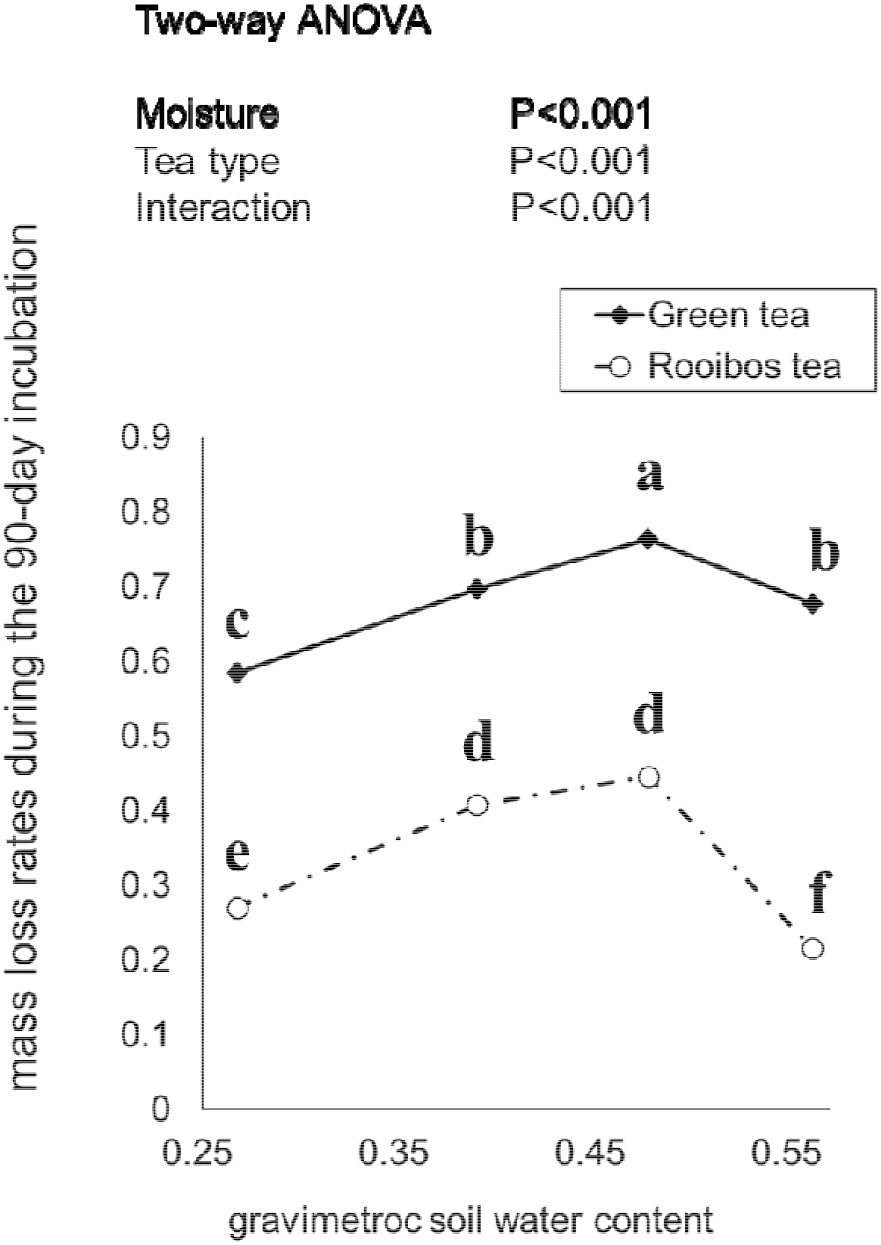
**Effects of water content on tea bag decomposition (Water Content Experiment). Tea bags were incubated with soils taken from a Japanese cedar plantation at four different water contents (27%, 39%, 48%, and 56%, wet weight basis). The error bars indicate the standard error of the four replicates**.

### Leaching loss of teas

Both tea type and temperature influenced the minimum leaching loss, with greater leaching for green tea and higher temperatures (Fig. 2a). Leaching loss when tea bags were submerged in 300 mL of water increased until 140 min, but was not further elevated at 24 h (Fig. 3) with the exception of green tea at 15°C (increasing until 24 h; Tukey’s post hoc test, Fig. 3). The results indicated that the leaching loss at 24 h was the maximum leaching loss. Tukey’s HSD indicated that (i) green tea experienced more leaching loss than rooibos tea and (ii) differences in temperature only affected the leaching loss from green tea at 10 min, where higher temperature caused a larger amount of leaching loss (Fig. 3).

**Fig. 2.**
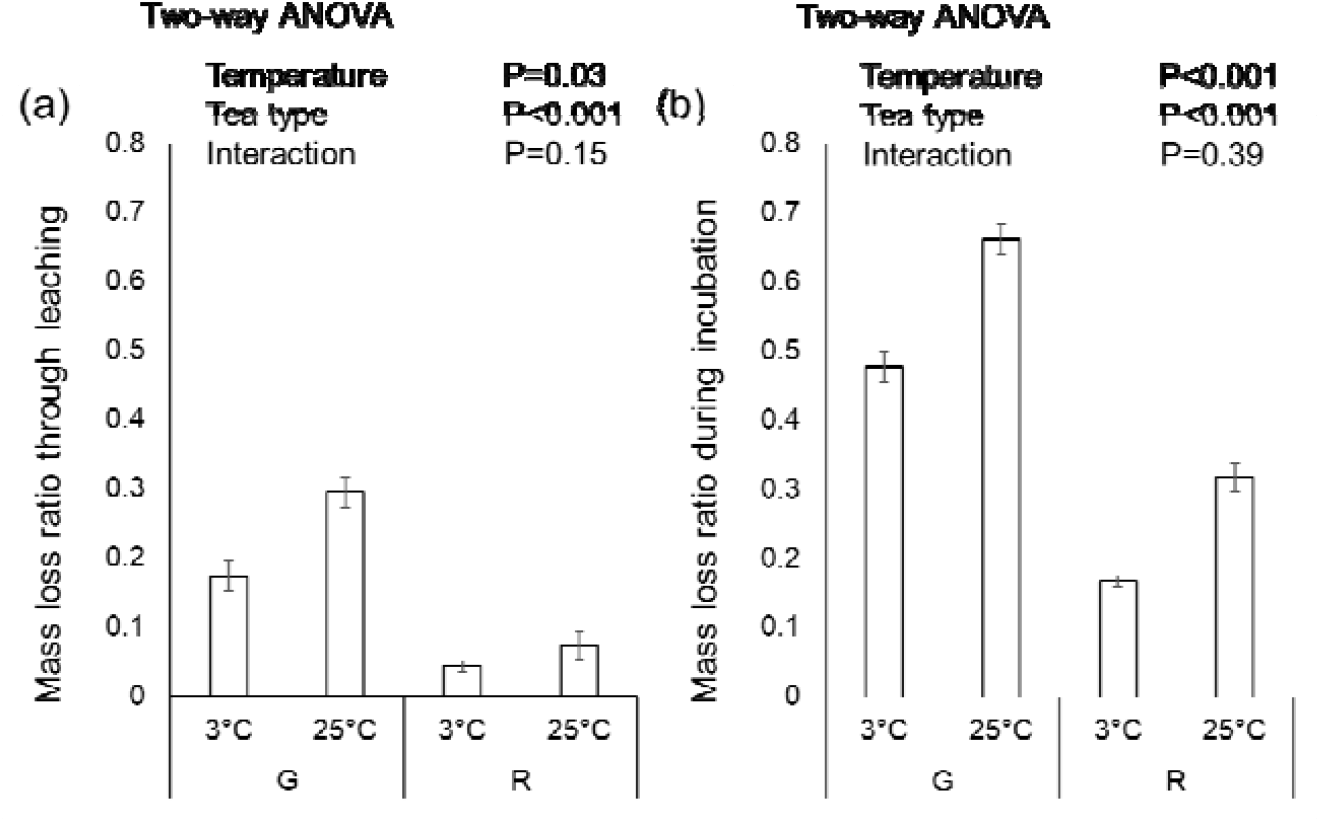
**Mass loss of green and rooibos teas relative to the initial weights (a) through leaching (Minimum Leaching Experiment) and (b) during a 90-day laboratory incubation period (Laboratory Decomposition Experiment). New tea bags made from polypropylene fabric (see main text) and soil taken from a Japanese cedar plantation were placed in PET bottles with drainage holes (see Fig. S3), and leaching and incubation experiments were conducted at 3°C and 25°C. The soil used for the leaching experiments was autoclaved at 120°C for 1 h to prevent any impact of microorganisms on mass loss of the tea bags. G, green tea. R, rooibos tea. The error bars indicate the standard error of the four replicates**.

**Fig. 3.**
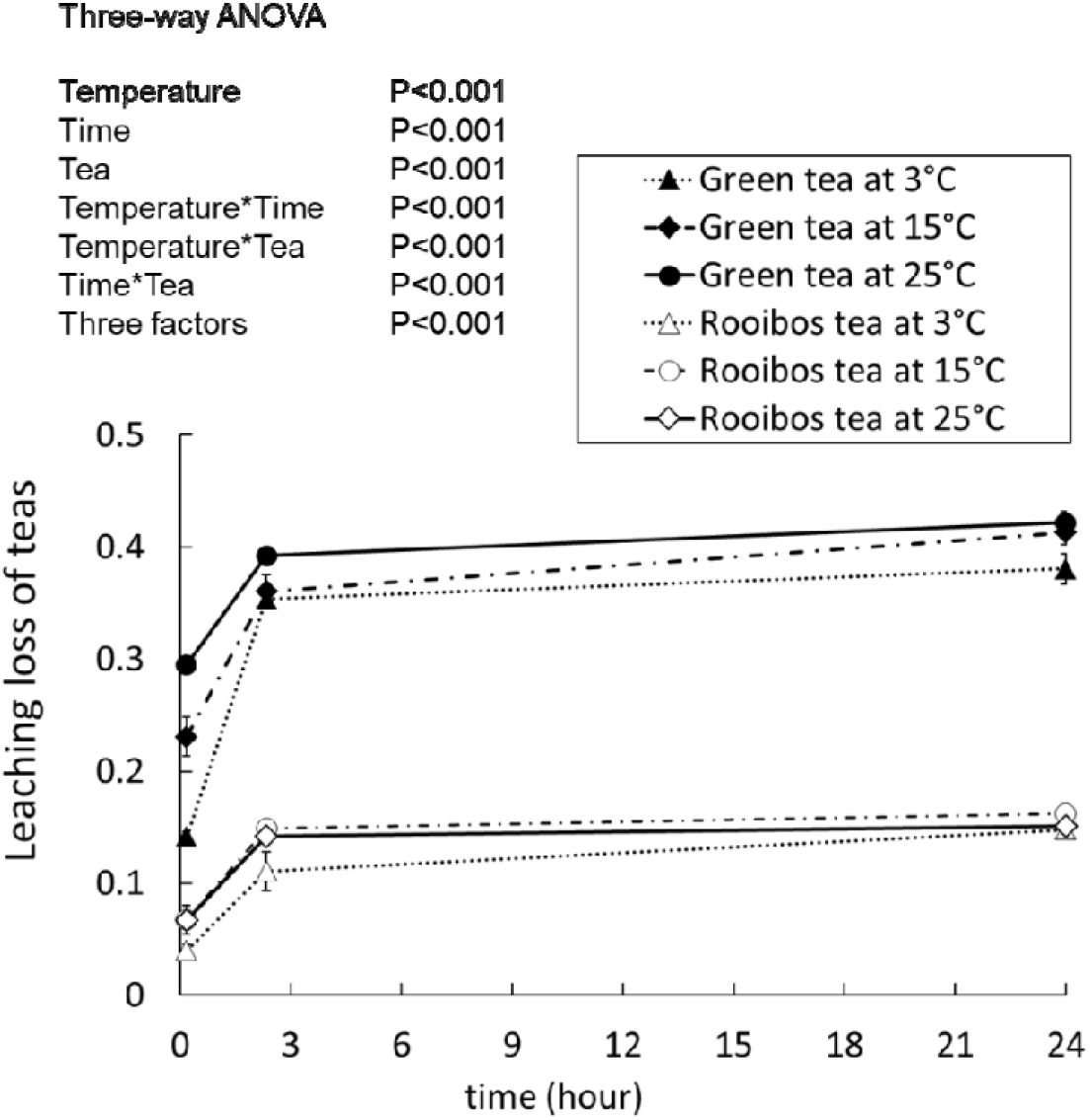
**Mass loss of green and rooibos teas relative to the initial weights during the Maximum Leaching Experiment. Effects of temperature and submergence duration on the ratio of leaching loss to the initial amount of green tea and rooibos tea (Maximum Leaching Experiment) were determined. New tea bags made from polypropylene fabric (see main text) were submerged in 300 mL water for 10 min, 140 min, and 24 h at 3°C, 15°C, and 25°C. The error bars indicate the standard error of the three replicates**.

### Contribution of leaching loss to tea bag decomposition

The lower bounds of the contribution of leaching loss to the tea bag decomposition rates ranged from 0.23 to 0.45, while the upper bounds reached as high as 0.89 (rooibos tea at 3°C) (Fig. 4). For green tea, the contribution of leaching loss to tea bag decomposition was largest at 25°C (Fig. 4).

**Fig. 4.**
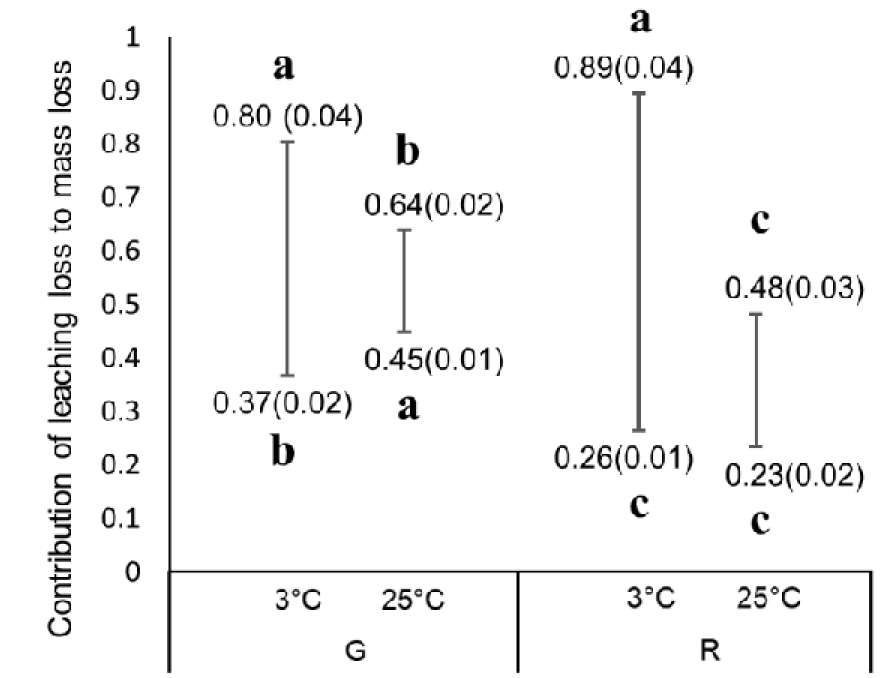
**Potential contribution of leaching loss to the mass loss of green and rooibos teas. Upper bounds were calculated as the ratio of the maximum leaching loss (Maximum Leaching Experiment) to the mass loss during the laboratory decomposition experiment (Total Mass Loss Experiment). Lower bounds were calculated as the ratio of the minimum leaching loss (Maximum Leaching Experiment) to the mass loss during the laboratory decomposition experiment (Total Mass Loss Experiment). Different letters on the graph indicate significant differences among the data for the upper bounds and lower bounds, determined using Tukey’s HSD followed by one-way ANOVA (P < 0**.**05). G, green tea. R, rooibos tea. The values on the graph are averaged data (with standard error) of the four replicates in parentheses**.

### Effects of mesh size on tea bag decomposition

The decomposition rate of green tea was not significantly affected by mesh size in either forest (Fig. 5), while the mass loss of rooibos tea (0.281 ± 0.01) in nonwoven bags was slightly but significantly higher than that of 0.25 mm-mesh bags (0.242 ± 0.01) at the Japanese cedar plantation (Fig. 5b).

**Fig. 5.**
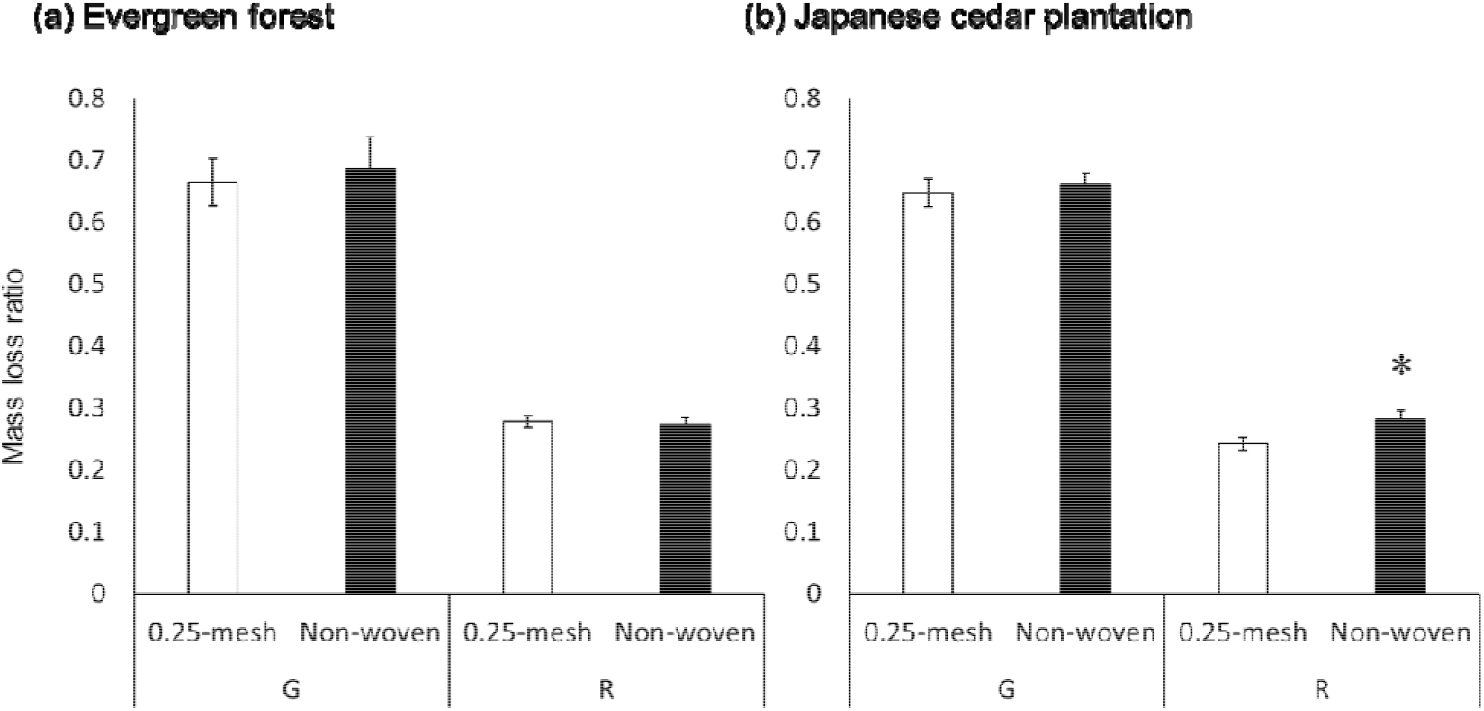
**Effects of mesh size on mass loss relative to the initial weight in (a) a mixed forest and (b) a Japanese cedar plantation. G, green tea. R, rooibos tea. 0**.**25-mesh, 0**.**25-mm mesh woven tea bags filled with tea from nonwoven teabags (Fig. S2). Nonwoven, polypropylene nonwoven tea bags with a non-uniform mesh size finer than 0**.**25 mm (see Fig. S1). *P < 0**.**05, old vs. new tea bags, paired *t*-test. Error bars indicate the standard error of the four replicates. Field incubation was performed for 90 days**.

## Discussion

### Effects of water content on tea bag decomposition

Both green tea and rooibos tea exhibited higher decomposition rates with higher water contents, but the highest water content inhibited decomposition (Fig. 1); this was consistent with the conceptual models of Prescott (2010). This result was most likely due to the reductive condition suppressing decomposition (Neckles & Neill 1994). The fact that the responses of both teas to the wetter condition were consistent with a conceptual model justifies the usage of tea bags to natural litters in studies of organic matter decomposition in wet conditions. Furthermore, the present results could help improve models of tea bag decomposition rates (ex. Didion et al. 2016). The “concave down shape” relationship between water content and tea bag decomposition rate could be adopted in such models.

### Potential contribution of leaching to tea bag decomposition rate

The leaching losses of rooibos tea after 24 h at 3°C, 15°C, and 25°C (0.15 ± 0.005, 0.16 ± 0.001, and 0.15 ± 0.003, respectively, Fig. 2) were comparable with those of other litters reported in previous studies (Nykvist 1959, Taylor & Parkinson 1988, Ibrahima et al. 2008). Ibrahima et al. (2008) reported the mass losses of leaf litters of the following eight agroforestry species after submergence in water: *Lophira lanceolata, Vitex doniana, Vitellaria paradoxa, Syzygium guineense* var. *guineense, Annona senegalensis, Syzygium guineense* var. *macrocarpum, Vitex madiensis*, and *Ximenia americana*. They demonstrated that all of the litters showed leaching losses below 20% during the 24-h submersion (all less than 10% except for *V. madiensis* and *X. americana*).

By contrast, green teas exhibited extremely large leaching losses (Fig. 1a). The leaching losses of green teas after 24 h at 3°C, 15°C, and 25°C (0.38 ± 0.01, 0.41 ± 0.01, and 0.42 ± 0.01, respectively) were much larger than in previous reports (Nykvist 1959, Taylor & Parkinson 1988, Ibrahima et al. 2008). The much larger fractions of nonpolar extractives and water solubles (hot water-extracted) in green tea compared to rooibos tea (Keuskamp et al. 2013) likely caused the larger leaching loss of green tea. The contrasting leaching losses of the two types of teas caused the larger potential contribution of leaching loss to the decomposition rate of green tea compared to that of rooibos tea, especially at the higher temperature of 25°C (Fig 3).

### Effects of mesh size on tea bag decomposition

Contrary to our hypothesis, the finer mesh size did not suppress the decomposition of rooibos tea. In the mixed forest, no significant difference in decomposition rate was observed between nonwoven and 0.25 mm-mesh tea bags, while in the cedar plantation the decomposition rate of rooibos tea was somewhat quicker for the finer nonwoven tea bags (although the mesh size effect was small; Fig. 5b). These results contrasted with earlier works, wherein finer mesh size caused slower decomposition (Bradford et al. 2002, Powers et al. 2009). It seems unlikely that the soil contamination caused the slower decomposition of the bags with coarser mesh, because the contamination was negligible in our study sites (personal observation). We cannot fully explain the phenomenon, which differed from the literature. One possible mechanism is as follows: a greater abundance of microorganisms decomposed litters due to the decrease in predation by mesofauna in the finer nonwoven bags. This hypothesis requires further study and quantification of mesofauna. On the other hand, green tea decomposition was not affected by the difference in mesh size, consistent with our initial hypothesis. Keuskamp et al. (2013) suggested that the decomposable fraction of green tea is completely decomposed before the end of the 90-day incubation period. In such a case, changes in mesh size and mesofauna abundance would not stimulate green tea decomposition.

Overall, the present study demonstrated that tea bag decomposition during a 90-day incubation period was not markedly affected by mesh size. Although the decomposition of rooibos tea was slightly stimulated by a change in the mesh size at the cedar plantation (from 0.242 ± 0.01 [nonwoven bags] to 0.281 ± 0.01 [0.25 mm-mesh bags], Fig. 5b), the impact was not large. Our unexpected results may justify combining the decomposition data of the two types of tea bags, namely, the previously produced woven nylon mesh tea bags and the currently used polypropylene tea bags. However, the number of observations in the present experiment was not sufficient to draw a definitive conclusion; additional studies with more observations are therefore necessary.

## Conclusion

The present study provides essential information for future studies on tea bag decomposition. First, we confirmed that both green and rooibos teas exhibited higher decomposition rates with higher water contents, although decomposition was inhibited at the highest water content; this was consistent with conceptual models of natural litters and justifies the assumption that tea bags are representative of natural litters. Second, we unexpectedly found that the impact of tea bag mesh size on decomposition might be insignificant. Thus, it may be possible to combine the decomposition data of the two types of tea bags, namely, the currently used bags and the previous ones, which are currently unavailable. Further studies with more study sites are required to test this.

## Acknowledgement

We thank all staff members of the Kamigamo experimental station, Field Science Education and Research Center, Kyoto University. We thank Dr. Yoshihiro Takahata for his professional advices on the laboratory work. We thank Ms Yumiko Sakamoto and Ms Akane Sakumori for their assistance on the laboratory work. This study was financially supported by JSPS KAKENHI Grant Number JP19K15879.

## Supporting information

**Fig. S1.**
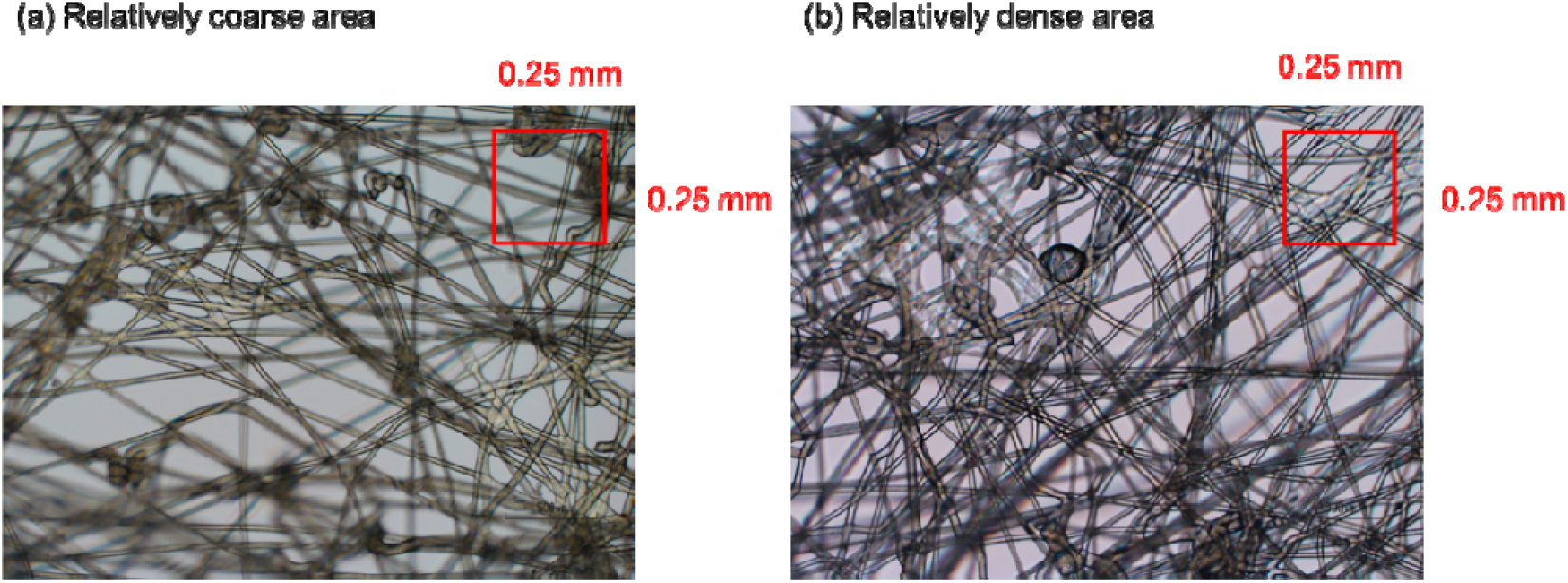
**Polypropylene nonwoven new bags with non-uniform mesh sizes. A relatively coarse area (a) and relatively dense area (b) are shown**.

**Fig. S2.**
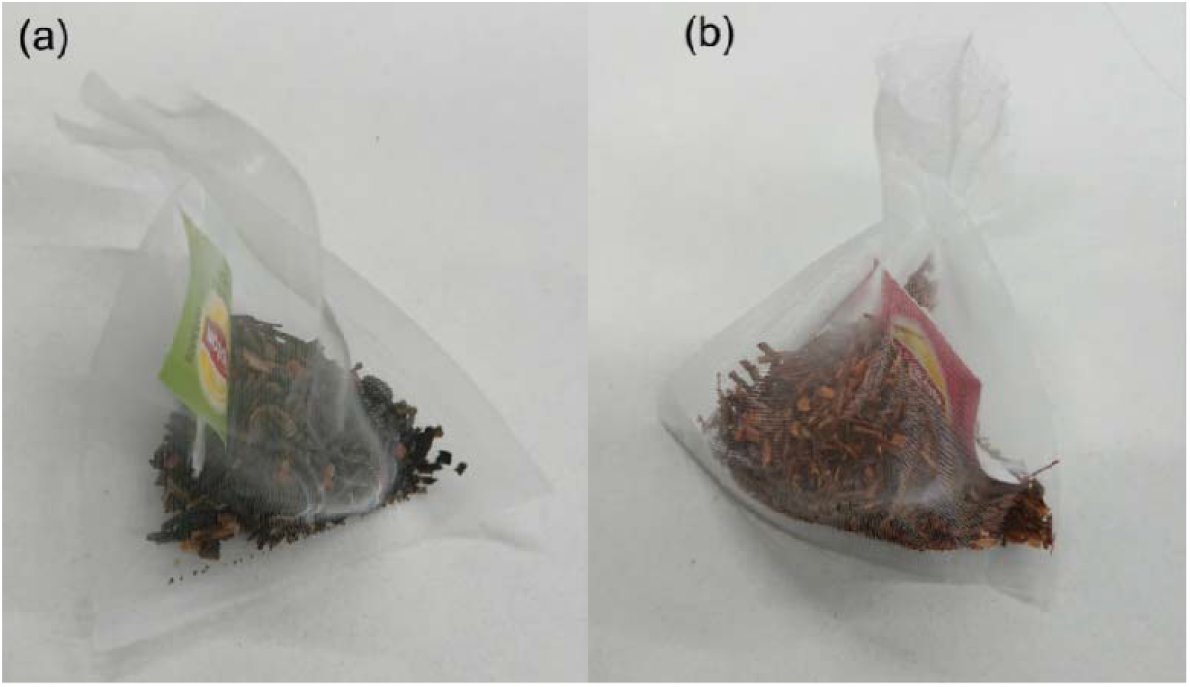
**Tea bags for (a) green tea and (b) rooibos tea made from 0**.**25-mm mesh and tea contained within the nonwoven tea bags**.

**Fig. S3.**
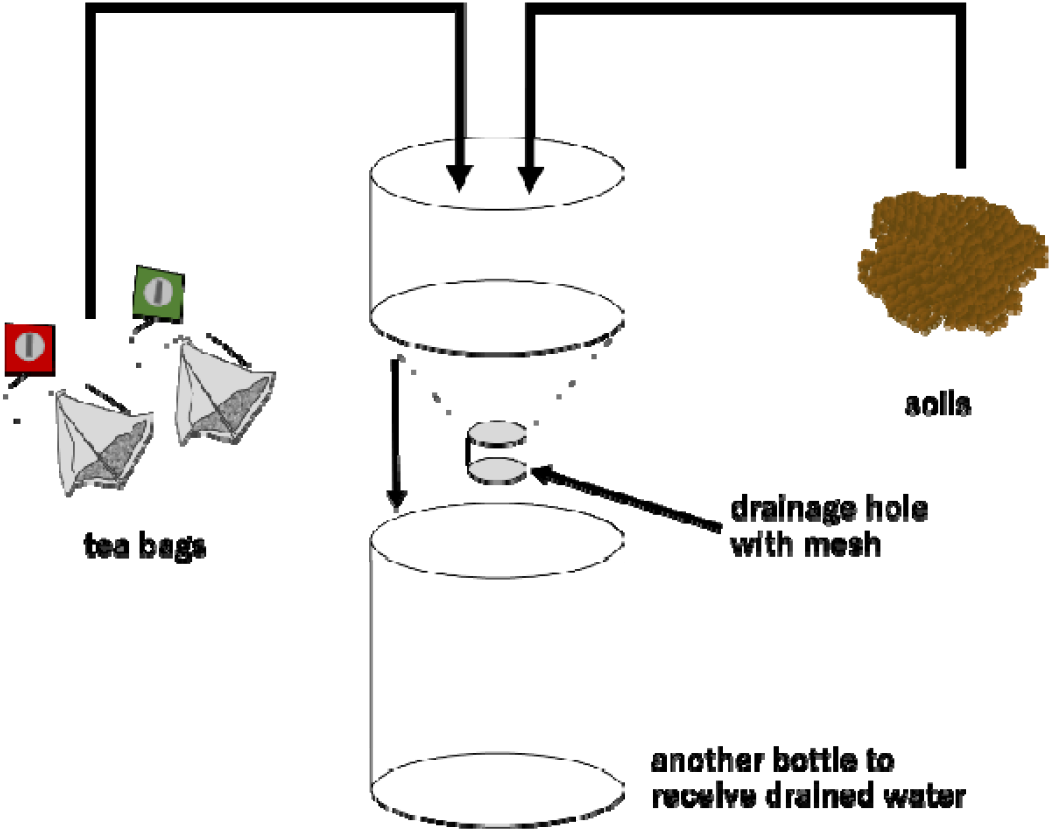
**Bottles used in the Total Mass Loss Experiment and Minimum Leaching Experiment.**

**Fig. S4. Climate data obtained during the field incubation period. Daily temperatures of the (a) evergreen forest and (b) Japanese cedar plantation, and daily precipitation amounts for the (c) evergreen forest and (d) Japanese cedar plantation, were determined by The Agro-Meteorological Grid Square Data, NARO.**

